# The bacterial lectin LecA from *P. aeruginosa* alters membrane organization by dispersing ordered domains

**DOI:** 10.1101/2022.04.17.488572

**Authors:** Taras Sych, Ramin Omidvar, Rafael Ostmann, Thomas Schubert, Annette Brandel, Ludovic Richert, Yves Mely, Josef Madl, Winfried Römer

## Abstract

The assembly and dynamic reorganization of plasma membrane nanodomains (also known as “lipid rafts”) play key roles in host cell infection by human pathogens (e.g. viruses and bacteria). Viruses and bacteria can trigger the reorganization of lipid rafts which leads to membrane invaginations and downstream signaling that promote infection. Such reorganizations can be induced by interactions of bacterial or viral carbohydrate proteins (so-called lectins) with lipid raft glycosphingolipids (GSLs). Here, we studied the GSL globotriaosylceramide (Gb3) which is a key receptor involved in the cellular uptake of the gram-negative bacterium *P. aeruginosa*. The bacterial surface lectin LecA targets Gb3 and promotes bacterial invasion via the “lipid zipper” mechanism. However, the impact of LecA on the organization of membrane nanodomains is unknown yet. We mimicked of the plasma membrane using supported lipid bilayers (SLBs) that contained liquid-ordered (Lo, “raft-like”, enriched in sphingolipids and GSLs) and liquid-disordered (Ld, “non-raft-like” enriched in DOPC) lipid domains. Upon interaction with LecA, the Lo domains in the SLBs reshaped and dispersed. Moreover, deformation of SLBs was observed as LecA formed membrane multilayers on SLBs surface. We further dissected this process to reveal the impact of Gb3 structure, bilayer composition and LecA valence on the Lo reorganization.

## Introduction

Lipid rafts are highly dynamic nanometer-sized plasma membrane domains enriched in sphingolipids, cholesterol and membrane proteins^1,2^. They play a key role in signal transduction^3–5^ and membrane deformation^6–8^ induced by human pathogens (viruses, bacteria) and pathogenic products (such as toxins) during the first steps of the infection of the host cell. Pathogens can target various plasma membrane components that can be found in lipid rafts, including glycosphingolipids (GSLs)^9–13^. The GSLs can be recognized by carbohydrate binding proteins (so-called lectins) that are often present on the surface of viruses and bacteria, or can be secreted by the latter. The lectin-GSL interactions can initiate signaling cascades^14^ or trigger membrane deformations^15,16^ by local remodeling of membrane content and organization^17–20^. Thus, a thorough study of the lectin-induced reorganization of membrane domains should provide valuable insights into the mechanisms of the initial steps of infection.

Due to their small size and highly dynamic character, it is still extremely challenging to characterize lipid rafts in living cells^21^. Therefore, less complex synthetic membrane systems^22^ with phase separation into Liquid-disordered (Ld) domains that mainly consist of phospholipids with unsaturated fatty acyl chains (e.g. DOPC) and Liquid-ordered (Lo) domains, which are enriched in sphingomyelin and cholesterol, are frequently used to mimic membrane heterogeneity^2,23^. The micrometer-scale of Lo domains makes it possible to study the reorganization of these lipid raft-like domains with common microscopy techniques. It is also possible to spike these systems with GSLs to study their impact on membrane organization as a result of GSL-lectin interactions.

In this work, we focused on the interactions of the lectin LecA from the Gram-negative bacterium *P. aeruginosa* with the GSL globotriaosylceramide (Gb3, also known as CD77 or P^k^ blood group antigen^24^). Gb3 has been described as an interaction partner of LecA^14,25–27^, which is crucial for host cell adhesion and internalization of *P. aeruginosa* through the so-called lipid zipper mechanism^11^. Gb3 is also well-known as the main receptor of the B-subunit of Shiga toxin (StxB) from *S. dysenteriae* ^28^. Both lectins share the same receptor but induce different signalling events^14^ and follow distinct intracellular trafficking pathways^26^. These differences may originate from the ability of the two ligands to induce different modifications of the local plasma membrane heterogeneity. The impact of StxB on membrane organization was thoroughly characterized using phase-separated synthetic membranes^6,29,30^. In particular, Windschiegl et al. showed that StxB shrinks lipid raft-like Lo domains in planar supported lipid bilayers (SLBs)^6^. Yet, the impact of LecA-Gb3 interactions on the organization of Lo domains is largely unknown. To explore the LecA-induced modifications of the Lo domains, we monitored the effect of LecA on phase-separated, Gb3-containing SLBs by fluorescence microscopy and atomic force microscopy (AFM). Moreover, we investigated the role of Gb3 structure and LecA valence in this process.

## Materials and methods

### Lipids, glycolipids and fluorescent lipophilic probes

Lipids (1,2-dioleoyl-sn-glycero-3-phosphocholine (DOPC), sphingomyelin (SM) and cholesterol (chol)) were purchased from Avanti Polar Lipids. The Gb3 “mixture” (ceramide trihexoside from porcine brain, consisting as a mixture of different fatty acyl chains varying in length, hydroxylation and degree of saturation) was from Matreya LLC. The Gb3-FSL^31^ (function-spacer-lipid) linked to DOPE was from Sigma Aldrich. The lipophilic membrane probe β-BODIPY C12-HPC (2-(4,4-Difluoro-5-Methyl-4-Bora-3a,4a-Diaza-s-Indacene-3-Dodecanoyl)-1-Hexadecanoyl-sn-Glycero-3-Phosphocholine), from now on referred to as HPC-Bodipy, was from ThermoFisher Scientific. The environment sensitive membrane marker Nile Red 12 S (NR12S)^32^ was provided by Dr. Andrey Klymchenko (University of Strasbourg, France).

### Lectins and lectin labeling

The pentameric B-subunit of Shiga toxin (StxB) was purchased from Sigma Aldrich. Recombinant LecA was produced from *Escherichia coli* following previously published procedures^33^. The recombinant prokaryotic lectin *α*Gal (RPL *α*Gal, from now on referred to as di-LecA (di-valent)) was from Glycoselect. Lectins were labeled with Cy5 mono-reactive dye (NHS ester) from GE Healthcare. Briefly, 1 μL of a 10 mg/mL solution of the amino-reactive probe dissolved in DMSO was added to 100 μL of a 1 mg/mL protein solution in PBS (^-^/^-^, Gibco) supplemented with 100 μM NaHCO_3_, pH 8.5. The mixture was incubated for 1 h at room temperature (RT) under continuous shaking. The labeled lectins were purified using Zeba Spin Desalting Columns (0.5 mL, MW cut-off: 7.0 kDa) from ThermoFisher Scientific.

### Preparation of solid supported lipid bilayers

The SLBs were established following the formerly published procedure^34^. Briefly, lipid mixtures were prepared in chloroform. For phase-separated SLBs a mixture of DOPC/chol/SM/Gb3 with a molar ratio of 37.5/20/37.5/5 and for non-phase-separated SLBs a mixture of DOPC/chol/Gb3 with a molar ratio of 65/30/5 were used. For fluorescence microscopy studies, SLBs were labeled by adding 0.1 mol-% of HPC-Bodipy to the lipid mixture in chloroform. Chloroform was removed by evaporation under nitrogen atmosphere and followed by vacuum (10 −15 mBar) application. Lipids were resuspended in ultrapure water at 55 °C with extensive vortexing in order to form large multilamellar vesicles. Then, the vesicle suspension was extruded using the LiposoFast-Basic extruder (Avestin) with 50 nm pore size filter (55 °C, 51 passages). A thin freshly cleaved layer of mica (PLANO, V1) was attached to a microscopy glass coverslip (Menzel Gläser, 30mm, #1.5) using UV-curing glue (Norland optical adhesive 63). The extruded small unilamellar vesicle (SUV) suspension (50 μL) was applied to the mica together with 150 μL of high (10 mM) CaCl_2_-solution in pure water. The coverslips were incubated for 30 min at 55 °C in order to form the non-phase-separated lipid bilayer. Thereafter, SLBs were washed with warm (55 °C) PBS buffer to remove lipids not attached to mica. Finally, SLBs were cooled down slowly (1-1.5 hours) to RT in order to induce lipid bilayer phase-separation on a microscopic scale. For membrane order studies, the environmental sensitive membrane marker NR12S (2 μL of 0.1 mM NR12S in DMSO) was added to the SLBs when the bilayer formation has already been completed. After 1-2 min of incubation, the lipid bilayer was washed mildly with PBS. Each SLB-containing imaging chamber was mounted on the microscopy dish holder (Interchangeable Coverglass Dish, Bioptechs) for imaging^22^. Lectins at a concentration of 200 nM in PBS were further applied to the SLBs in the imaging chamber.

### Fluorescence Correlation Spectroscopy

To assess the quality of SLB preparation, lipid mobility within SLBs was measured using Fluorescence Correlation Spectroscopy (FCS). For FCS measurements, the concentration of HPC Bodipy was reduced to 0.005 mol % for preparation. The correlation curves and diffusion coefficients are displayed at the Figure S8.

FCS measurements were carried out using Picoquant MT 200. 488 nm pulsed laser diode was used to excite HPC-Bodipy. 40X 1.2 NA water immersion objective was used to focus the light. 5 curves were taken per spot (5 s each). Laser power was set to 0.1-0.5 % of the total laser power that corresponds to 2-10 mW. Curves were then fitted using the SymphoTime software 2D-diffusion model below ^17^.

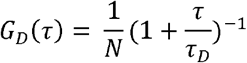

where N represents the number of fluorescent species within the beam’s focal volume. Next, the diffusion coefficients were calculated as follows:

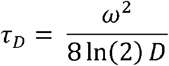

Where *w* corresponds to the full-width half maximum of the point spread function; *T_D_* is the diffusion time and *D* is diffusion coefficient.

### HILO-illumination microscopy

SLBs were often imaged using TIRF microscopy^23^. However, in our case, the specific design of the SLB support may restrict the total internal reflection of the laser beam at the mica/SLB interface. Therefore, we used an optical configuration similar to HILO^35^ – highly inclined thin layer illumination below the angle of total internal reflection. This method provides a high signal to background ratio decreasing the illumination of the non-bound lectins in solution. HILO-illumination imaging was performed with a Nikon Eclipse Ti-E fluorescence microscope (with 100x oil objective, N.A. 1.49 and an iXon DU-897 EMCCD camera (Andor Technology)). The 488 nm and 647 nm lasers were used to excite HPC-Bodipy and Cy5, respectively. After lectin application, several areas of each SLB were imaged with exposure times of 30-50 ms and intervals of 10-20 seconds between frames.

### Time lapse data analysis

The resulting time-lapse images of the SLBs were analyzed using a Fiji-based, home-made macro^36,37^. First, bleaching and laser-profile corrections were applied. Then, the total area of Lo domains, the surrounding Ld membrane and furthermore also membrane defects and multilayers were calculated for each frame. In addition, the sizes of the individual Lo domains were extracted. The macro is available at https://github.com/taras-sych/Tools-for-SLBs.

### Atomic Force Microscopy

The phase-separated SLBs were prepared slightly different as described above. Instead of passive cooling at RT, the small unilamellar vesicle suspension on the mica was placed in the Bruker JPK Biocell holder for 5-10 min. The holder was filled with warm PBS (55 °C) and subsequently washed thoroughly in order to remove unattached lipids. Using the JPK experiment planner module of the Bruker JPK SPM software, the cooling step was programmed in several steps (15 min at 55 °C and gradual temperature decrease to a temperature of 45 °C with 2 °C steps every 6 min and then decrease from 45 °C to the final temperature of 22 °C with 2 °C steps every 3 min). Afterwards, the holder was kept at the final temperature during the entire measurement. This stepwise cooling procedure resulted in uniform separation of lipid domains and prevented buffer evaporation.

The AFM imaging was performed using sharp nitride cantilevers (Bruker SNL-10 tip A) with a nominal tip radius of 2 nm. With the AC imaging mode, we imaged areas of 10 μm x 10 μm and 20 μm x 20 μm with the resolution of 512 x 512 pixels at a line scan rate of 2 Hz.

After the recording of some areas of the intact bilayer, we added diluted LecA solution to the Biocell holder (to have a final concentration of 100 nM LecA) and started to image the bilayer. Typically the first image was recorded around 2-3 mins afterLecA addition.

The editing of the images, such as plane subtraction to remove the image tilt and line replacements, was done using JPK data processing software. The line profiles and histograms were extracted from JPK data processing software and plotted using Inkscape.

### Confocal ratiometric microscopy

The two-color ratiometric imaging was performed on a confocal fluorescence microscope (Nikon Eclipse Ti-E with A1R confocal laser scanner, 60x oil objective, N.A. 1.49). For fluorescence excitation, the 488 nm laser was used. The emission of NR12S was simultaneously recorded in two color channels - green (using 525/50 BP filter) and far red (700/75 BP filter).

The two-color-channel raw image was analyzed using a home-made Fiji-based macro. First, the threshold was applied to remove signal from pixels that represented membrane defects. Second, in order to calculate the general polarization^38^ (GP) value of the NR12S spectrum for each pixel of the image the following formula was used:

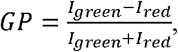

where *I_green_* is the gray value of the pixel in the green color channel and *I_red_* is the gray value of the pixel in the red color channel. The resulting GP image contained GP values for each pixel of the image. The GP histogram was extracted from the resulting GP image. NR12S is sensitive to membrane hydration and exhibits a drastic red shift in polar environments (i.e. in Ld domains). The GP value of NR12S informs about the local membrane order and can vary from −1 to 1. Lower values correspond to low membrane order whereas higher GP values indicate a higher membrane order. The Fiji macro is available at https://github.com/taras-sych/General-polarization.

### Statistical data analysis and representation

Data are represented using box-whiskers plots, where the middle horizontal line represents the mean value, the boxes the 25^th^ to 75^th^ percentiles, and the whiskers the standard deviation. The statistical significance was analyzed with Kruskal-Wallis non-parametric test due to non-normal distributions in all datasets.

## Results

### LecA disperses Lo domains and forms membrane multilayers

The heterogeneity of the outer leaflet of the plasma membrane was mimicked using phase-separated SLBs^22^. The phase-separation was visualized using the fluorescent label HPC-Bodipy, which preferentially incorporated into the Ld domains, whereas the Lo domains were depleted of HPC-Bodipy and appeared dark (figure 1a - e.g. blue arrowhead at 0 min). Importantly, Lo domains were stable over time in absence of LecA (Figure S2). Furthermore, membrane defects, which occasionally formed spontaneously during SLB preparation were stable over time, they neither shrank nor grew. Moreover, the quality of SLBs was tested by applying the B subunit of Shiga toxin (StxB), which binds specifically to Gb3. StxB is known to recognize Gb3 in Lo domains, reshape existing Lo domains and induce formation of new Lo domains by clustering Gb3 receptor molecules^6,30^. Importantly, we observed similar effects of StxB on our SLB experiments: StxB bound Gb3 in Lo domains and induced the formation of new Lo domains (Figure S3). These two experiments (Figures S2 and S3) can be considered respectively, as “negative” and “positive” controls, which validate the applicability of our SLBs for studying of lectin-induced membrane reorganization.

**Figure 1.**
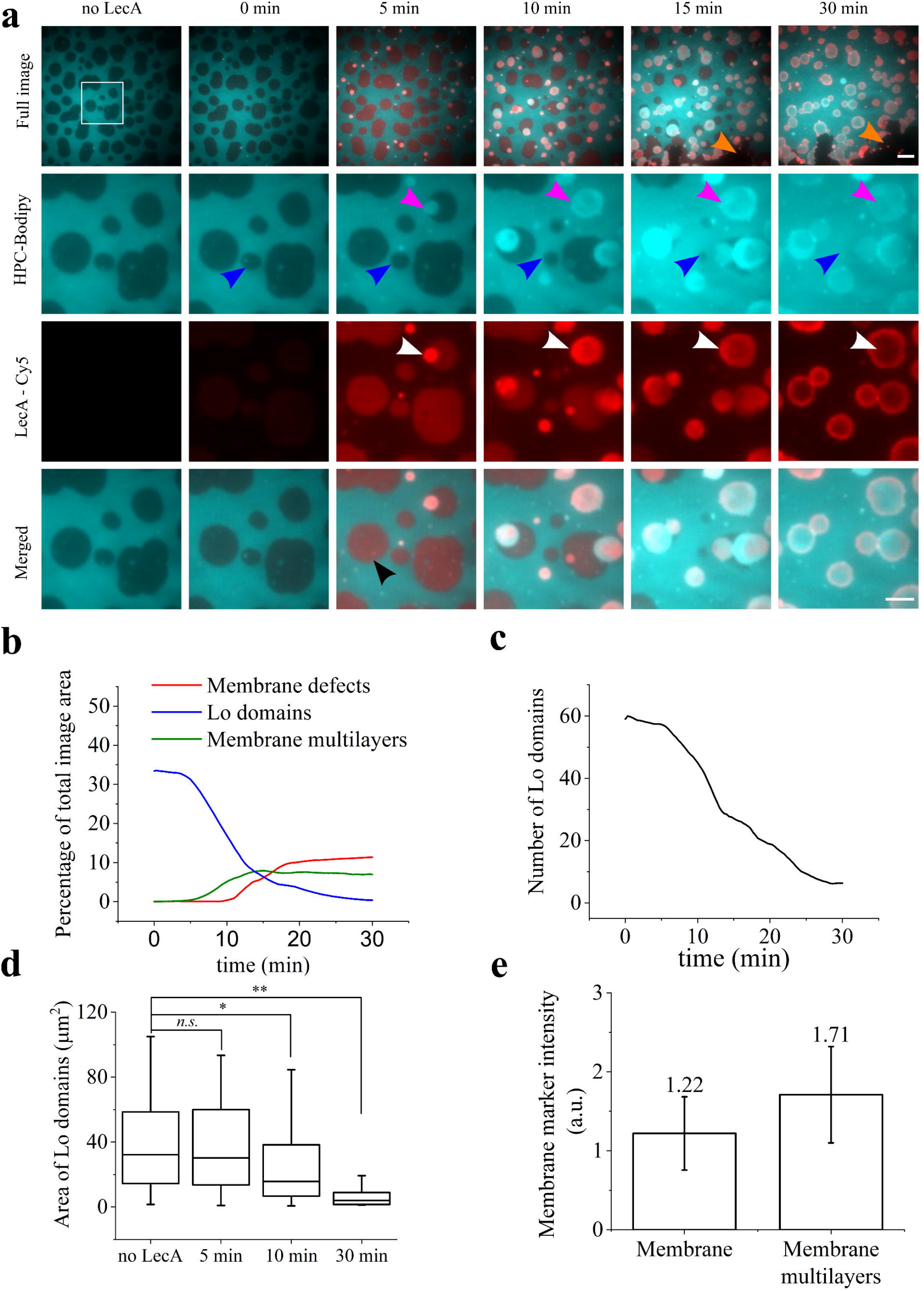
LecA-induced SLB reorganization visualized by fluorescence microscopy. The SLB was labeled by HPC-Bodipy that localized to Ld domains (cyan). LecA was labeled with Cy5 (red). **a)** Time series of LecA interaction with a phase-separated SLB (DOPC/chol/SM/Gb3 (37.5/20/37.5/5)). LecA bound mainly to Lo domains (black arrowhead, as one example), induced the dispersion of Lo domains (blue arrowheads) and later the formation of membrane multilayers (pink arrowheads). At around 5 min, LecA started to form clusters (white arrowheads) that colocalized with membrane multilayers. At later time points of incubation, large membrane defects appeared (orange arrowheads). Photobleaching of the fluorophore Cy5 coupled to LecA in membrane multilayers can be observed at later time points (30 min – white arrowheads). The complete time sequence is available online as supplementary movie 1. Scale bars are 10 μm (for full image) and 5 μm (for zoom). **b)** Changes of the total area of Lo domains, membrane defects and membrane multilayers over time. **c)** Changes of the total number of Lo domains over time. **d)** Changes of the sizes of the individual Lo domains over time. **e)** The mean fluorescence intensity in membrane multilayers (recorded at 15 min after LecA application) was approximately 1.5 times higher than the intensity of the original, single bilayer membrane.

Cy5-labeled LecA was incubated with SLBs at a concentration of 200 nM. LecA (presented in red color) bound mainly to Gb3 molecules in Lo domains (figure 1a – 0 min and supplementary movie 1). Interestingly, over time, LecA induced the dispersion of Lo domains (figure 1a – the blue arrowhead points to one of many events). The total area of the Lo domains decreased, and after about 30 min of incubation with LecA, the Lo domains disappeared completely (blue curve in figure 1b). A more detailed analysis of the dynamics of Lo domain dispersion revealed a reduction of the number (figure 1c) and the area of individual Lo domains (figure 1d).

Approximatively, 5 min after application of LecA, intense LecA clusters emerged preferentially at the Lo/Ld phase domain boundaries (figure 1a – pink arrowheads and supplementary movie 1). These clusters grew over time to micrometer sizes. Interestingly, simultaneously with the appearance of these LecA clusters, the growth of novel membrane patches was observed (figure 1a). These patches co-localized with the LecA clusters, and HPC-Bodipy fluorescence intensity in these patches was approximately 1.5 times higher than in the Ld membrane (figure 1e). As published by Villringer et al., tetrameric LecA has the ability to crosslink two opposing lipid bilayers upon which it strongly accumulates at membrane interfaces^39^. Accordingly, we presume that these transformed membrane areas represent membrane-LecA-membrane “sandwiches”, comprising two or even multiple lipid bilayers cross-linked by LecA. Furthermore, after 15 min, large defects in the SLB started to appear (figure 1a – orange arrowheads and supplementary movie 1). The total area of membrane defects increased over time (figure 1b), paralleling the increasing area of membrane multilayers (figure 1b – red curve). Importantly, the membrane multilayer formation preceded the membrane destruction (green curve in figure 1b), suggesting that the formation of multilayers depleted the initial SLB in lipids, which resulted in extensive membrane defects.

### LecA forms membrane multilayers

The thickness of membrane multilayers was assessed at the nanoscale by atomic force microscopy (AFM). In comparison to fluorescence microscopy, AFM generally provides slower acquisition speed (every image took about four minutes with the settings mentioned above). To allow meaningful recordings with AFM, the LecA concentration was reduced to 100 nM in order to slow down LecA-induced membrane reorganization processes. Before the addition of LecA, Lo domains were approximately 1.5 nm higher than Ld domains (figure 2 – no LecA). At 10 min after application of LecA, LecA molecules were visibly bound to both Lo and Ld domains (figure 2 – 10 min). Interestingly, the height of LecA bound to Ld domains was about 2 nm whereas its height on Lo domains was between 2.5 and 3 nm. As the dimensions of LecA are 7 nm x 3 nm x 2 nm, we speculate that LecA positioned on the membrane “horizontally” (i.e. with its longest axis parallel to the SLB plane) or at least tilted. The differences in LecA heights on Lo and Ld domains can be explained by the fact that membrane order can alter the exposure of Gb3 head groups, which in turn affects LecA binding and localization on the lipid bilayer surface^40,41^. After 60 min, clusters of 8 nm in height can be observed on top of Lo domains (figure 2 – 60 min). These clusters are likely composed of the LecA in “vertical” position (i.e. with its longest (7 nm) axis perpendicular to the SLB plane). After 90 min membrane multilayers of 23 nm in height appear (figure 2 - 90 min). These membrane multilayers probably contain a second layer of membrane on top of the initial SLB with LecA sandwiched in between. Moreover, LecA can bind on top of the second layer, hence forming SLB – LecA – membrane – LecA multilayer which adds up to 23 nm in thickness.

**Figure 2.**
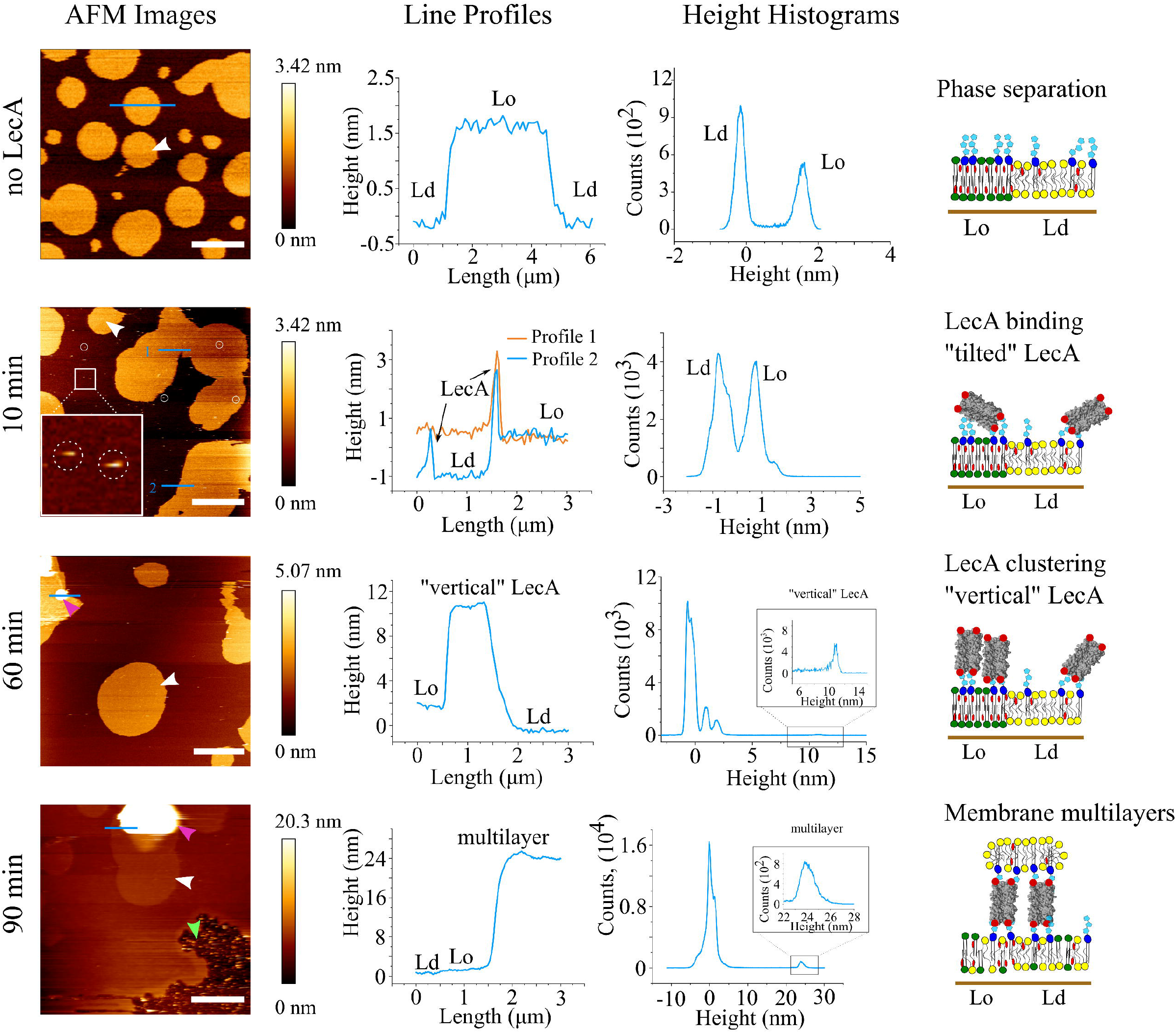
LecA-induced SLB reorganization visualized by AFM. Time series of LecA interaction with a phase-separated SLB (DOPC/chol/SM/Gb3 (37.5/20/37.5/5)). Time points (left panel) are collected from different locations at the SLB. **No LecA:** White arrowheads indicate Lo domains; **10 min:** White circles and white dashed circles (at zoomed area) highlight LecA bound to the SLB; **60 min:** Pink arrowhead indicates LecA cluster on top of Lo domains. **90 min:** Pink arrowhead indicates multilayer, green arrowhead points to an area of growing membrane defects. Scale bars 5 μm. The negative values are present due to differences in leveling between different pictures. Height profiles (middle left panel) were extracted along the blue lines as shown in the topography images. Height histograms are depicted in the middle right panel. The right panels show graphical representation of SLB and LecA structures presented in the images.

AFM imaging enabled us to confirm the formation of membrane multilayers with LecA sandwiched in between the lipid bilayers. However, the analysis of the lipid composition and order in membrane multilayers requires another approach, which is more sensitive to membrane environment in the lipid bilayer. In this work, we assessed this by studying the membrane order of multilayers using the environmental sensitive membrane probe NR12S.

### LecA drastically alters membrane order

Lo domains enriched in SM and chol exhibit a highly ordered lipid organization, while the surrounding Ld membrane environment is of low order^2,15^. After LecA-induced Lo domain dispersion, SM and chol are thought to redistribute and modify the membrane order accordingly. Various techniques provide the possibility to measure the local membrane order of a lipid bilayer^2,42^. In this work, we used the environmental sensitive membrane marker Nile Red 12 S (NR12S) to monitor the lipid order by fluorescence-based ratiometric imaging^32^.

We constructed the general polarization (GP) image to map the membrane order of the SLB (figure 3a). In absence of LecA, the GP image displayed a clear distinction between Lo domains (figure 3a – red/green, with a GP value of approximately 0.3) and Ld domains (figure 3a – blue, with a GP value of around −0.5). The two distinct populations that correspond to Lo and Ld domains are also clearly highlighted in the GP histogram (figure 3b – black curve). After application of 200 nM of LecA, the Lo domains dispersed, and the formation of multilayers and membrane disintegration were detected (figures 3a and S2), as described above. Similarly to the observations before, Lo domains shrunk and rearranged (figure 3a – white ROIs at 10 min and 18 min). The complete dispersion of the Lo domains was not observed in this experiment as membrane disintegration started very early (at 18 min time point), and SLB was completely disintegrated after 30 min (figure S2). In line with the observed decrease in the total area of Lo domains at 10 and 20 min of LecA incubation (figure S2), the Lo population of the GP histogram appears reduced accordingly, whereas the Ld population increased (figure 3b – red and blue curves).

**Figure 3.**
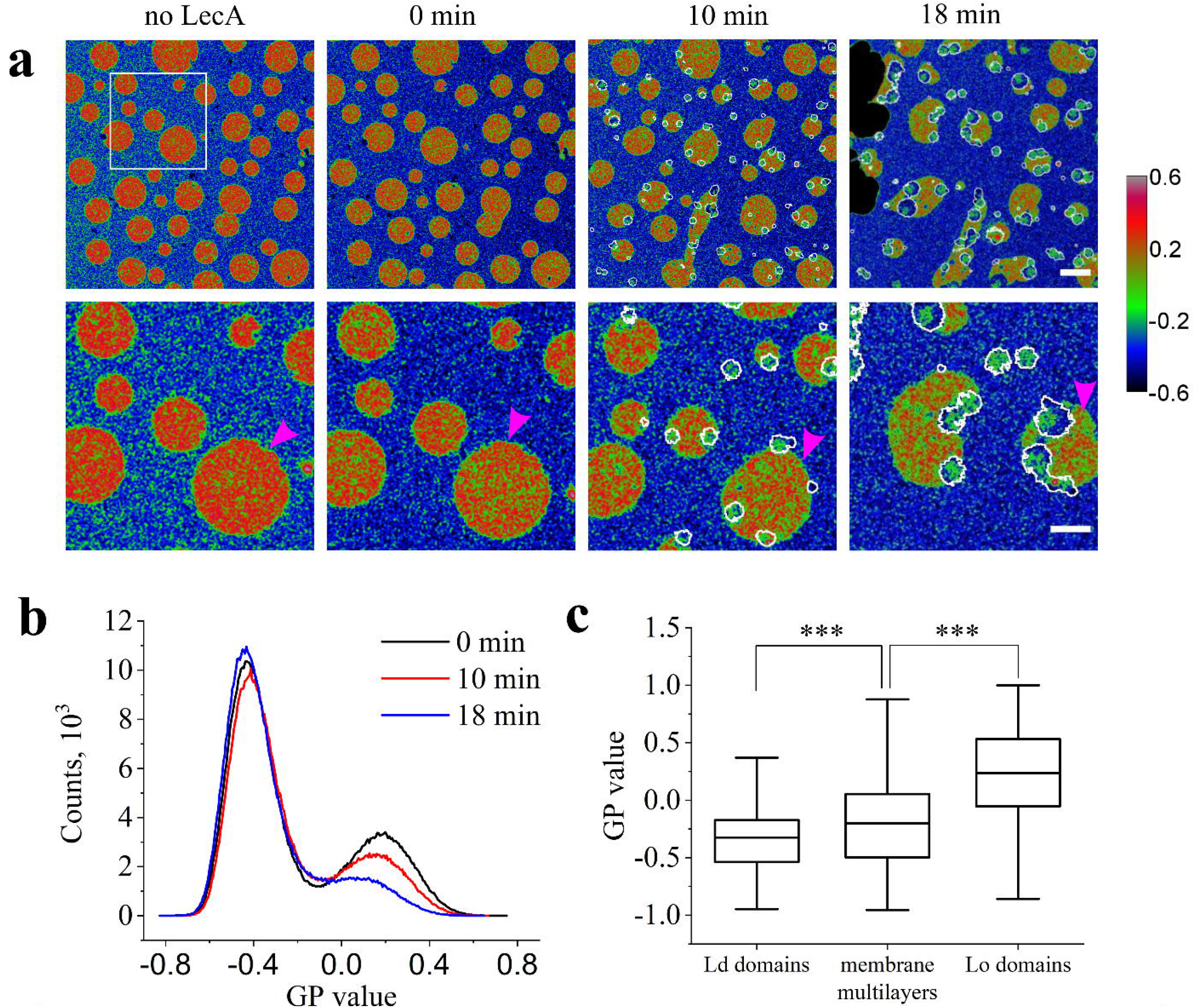
Membrane order changes in SLBs induced by LecA. Two-color imaging was performed using confocal microscopy. **a)** GP images of the SLB before and at several time points after LecA application. The color bar on the right panel encodes the GP values. Phase dispersion (pink arrowhead) occured in the same way as in figure 1. The membrane multilayers are highlighted by ROIs with white borderlines. **b)** GP histograms of the SLB at 0 min (black curve), 10 min (red curve) and 18 min (blue curve) after LecA application. **c)** Membrane order of Lo domains, Ld domains and membrane multilayers. Scale bar 10 μm. The ratiometric imaging is available in figure S2 and supplementary movie 2.

The formation of membrane multilayers was also observed in this experiment. We identified membrane multilayers using the raw two-color confocal images (figure S3). Again, the majority of membrane multilayers was observed at the Lo/Ld domain boundaries. In the resulting GP images, multilayers are highlighted by ROIs with white borderlines (figure 3a). These ROIs were used to quantify the GP in membrane multilayers. As a result, the membrane order of multilayers (−0.20 ± 0.36) was found to be higher than the order of Ld domains (−0.32 ± 0.36) but lower than the order of Lo domains (0.24 ± 0.37) of the original lipid bilayer (figure 3c).

This intermediate membrane order is likely the consequence of lipid redistribution in the membranes caused by the dispersion of Lo domains. However, another process, namely the attachment and growth of a Ld – like membrane on top of an existing Lo domain can produce the same result and explain the dispersion of Lo domains. To get further information, we wanted to determine whether the phase dispersion and membrane multilayer formation are synergetic or whether they can occur independently. In order to observe multilayer formation in absence of the dispersion of Lo domains, we designed two experiments. First, we produced phase separated SLBs supplemented with 5 mol % of synthetic construct Gb3-FSL-DOPE (further referred as Gb3-FSL) instead of the Gb3 wild type mixture, used above. The Gb3-FSL construct incorporates exclusively in Ld, hence LecA can only bind to the Ld domain of the SLB. In this case, membrane multilayer formation and SLB disintegration start within seconds after LecA application to such SLB (figure S5). Within minutes, the SLB was completely disintegrated. Interestingly, the dispersion of Lo domains was not observed and SLB patches constituted of Lo membrane could be found still attached to the SLB substrate at the late time points (Figure S5). In a second experiment, we produced homogeneous SLBs consisting of DOPC/chol/Gb3 (65/30/5), where phase separation does not occur, and thus, Lo domains cannot be dispersed. In this system, LecA (200 nM) induced multilayer formation and membrane disintegration (figure S6) very fast. Similarly to phase-separated SLBs with Gb3-FSL, the membrane was completely disintegrated within 15 min. These two experiments showed clearly that membrane multilayer formation can occur in the absence of Lo domains dispersion.

Next, we tested whether phase-dispersion is possible in the absence of membrane multilayer formation. For that we employed the homodimeric variant of LecA (di-LecA).

### The dimeric LecA variant disperses Lo domains without forming membrane multilayers

The di-LecA and wt-LecA monomers share the same amino acid sequence, with the exception of the N-terminus, which is modified with a 6xHis-tag in di-LecA monomers ^43^. This modification stabilizes di-LecA as a dimer in aqueous solution and does not affect its specificity to the *α*-galactose moiety of glycoconjugates (e.g. Gb3). Applied to phase-separated SLBs containing Gb3, di-LecA (200 nM) recognized preferentially Lo domains (figure 4a). Membrane reorganization was triggered immediately after di-LecA addition. Over the first 15 min, the total area of Lo domains decreased (figure 4a-c) as a result of di-LecA-induced fusion of Lo domains (figure 4b). At the 15 min time point, the reshaping of Lo domains stopped. Nevertheless, the ratio of the fluorescence intensity of HPC-Bodipy in Ld and Lo domains (I_Ld_/I_Lo_) indicates a two-fold drop over the next 45 min (figure 4 a, d, e). This might be explained by an “asymmetric” Lo domain dispersion. Namely, we hypothesize that in the upper membrane leaflet, which is directly exposed to di-LecA, Lo domains disperse, whereas in the lower membrane leaflet the complete dispersion of Lo domains is constrained by the mica substrate. Moreover, the binding of di-LecA at late time points (figure 4a – 60 min) became more homogeneous, suggesting an efficient mixing of the Gb3 molecules in the upper leaflet of the lipid bilayer. Interestingly, similar events were also observed in experiments with the tetrameric wt-LecA (supplementary movie 1), but, in this case, the events were very fast and resulted in the complete dispersion of Lo domains.

**Figure 4.**
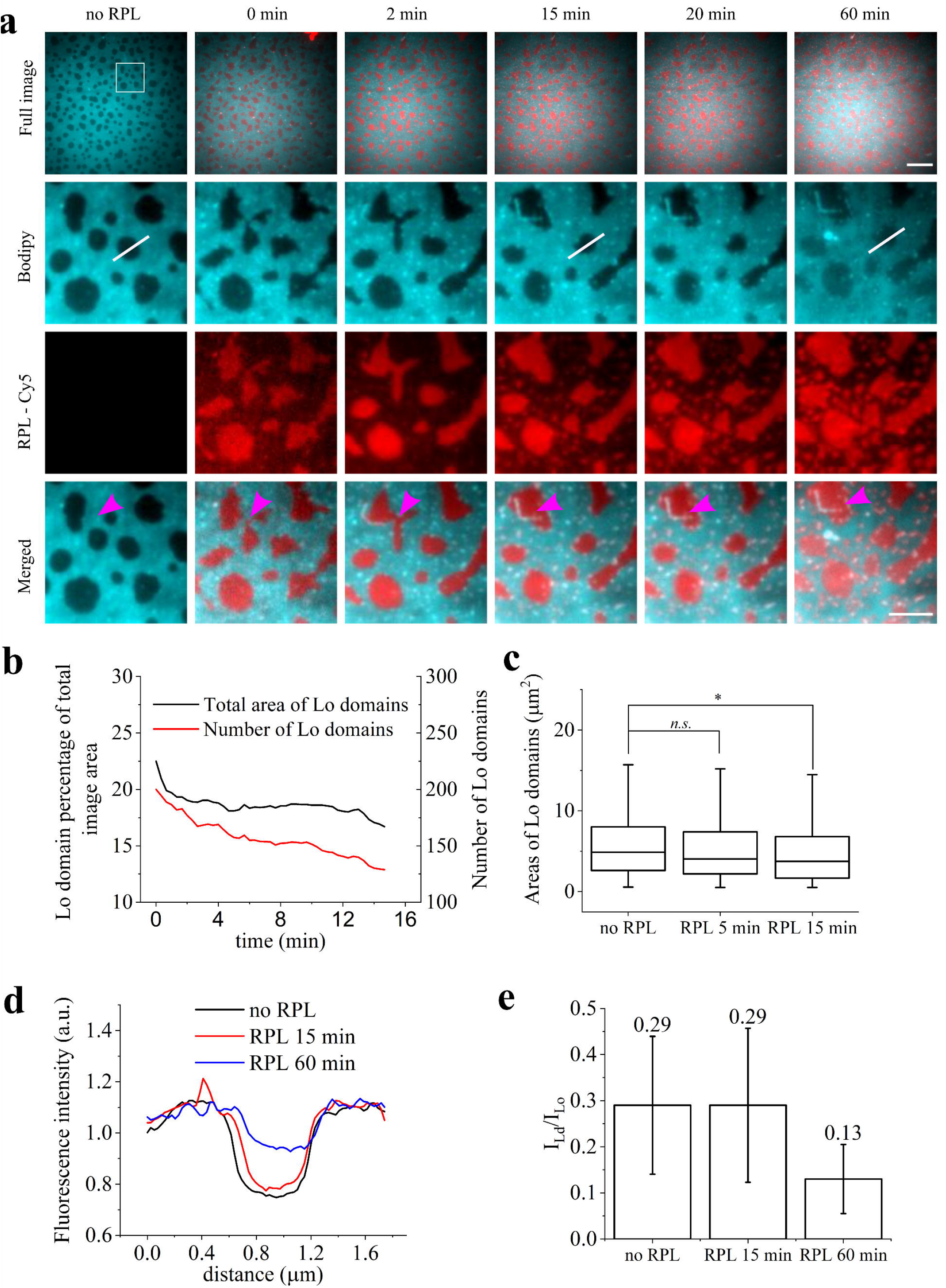
di-LecA-induced SLB reorganization visualized by fluorescence microscopy. The SLB was labeled by HPC-Bodipy that localized into Ld domains (cyan). di-LecA was labeled by Cy5 (red). Images were acquired with wide field microscopy (HILO). **a)** Time series of di-LecA interactions with phase-separated SLB (DOPC/chol/SM/Gb3 (37.5/20/37.5/5)). di-LecA bound preferentially to Lo domains and induced fusion and a decrease in size of the Lo domains (pink arrowhead). **b)** Decrease of the total area and number of Lo domains over time. **c)** The sizes of the individual Lo domains slightly decreased over time; **d, e)** Contrast of the fluorescence signal between the Ld and Lo domains at different time points. **d)** HPC-Bodipy fluorescence intensity profile along the white line displayed in (a) at different time points. **e)** Intensity contrast (ratio between the fluorescence intensity signals of HPC-Bodipy in Ld and Lo domains) value at different time points for all Lo domains. After 1 h, the contrast dropped to about half of the initial value. This suggests a so-called “asymmetric” Lo domain dispersion (i.e. Lo domains disperse only in the upper lipid monolayer whereas in the bottom monolayer, they remain intact). The scale bars are 10 and 5 μm, respectively. The complete time sequence is available online as the supplementary movie 4.

Application of di-LecA to SLBs with Gb3-FSL led to reshaping of Lo domains, but no Lo dispersion (neither asymmetric nor full) was observed (Figure S7). This confirms that the presence of receptor in the Lo domain is crucial for Lo dispersion induced by LecA.

Membrane multilayer formation was not observed in these experiments, clearly indicating that dispersion of the Lo domains can occur in the absence of membrane multilayer formation.

## Discussion

In this work, we demonstrated an impact of LecA on the (re-)organization of model membranes. LecA was found to induce the dispersion of Lo domains, multilayer formation and membrane disintegration on phase-separated SLBs. This effect of LecA is not a common feature of Gb3-binding lectins. Lectin-induced membrane reorganization processes were formerly studied in the context of the cellular uptake of the B-subunit of Shiga toxin (StxB) from *S. dysenteriae*. On synthetic membrane systems (GUVs^30^ and SLBs^6^), StxB bound Gb3 molecules in Lo domains. Moreover, StxB could reshape and fuse existing Lo domains (demonstrated by Windschiegl et al. ^6^) and induce formation of novel Lo domains (demonstrated by Safouane et al. ^30^). We reproduced these results in our SLBs as well. Both lectins (namely LecA and StxB) are specific to the same receptor but their impact on the organization of model membranes differs significantly. The reasoning behind such diversity as well as the proposed mechanism of LecA activity at the membrane are summarized and discussed below point by point as well as in figure 5.

**Figure 5.**
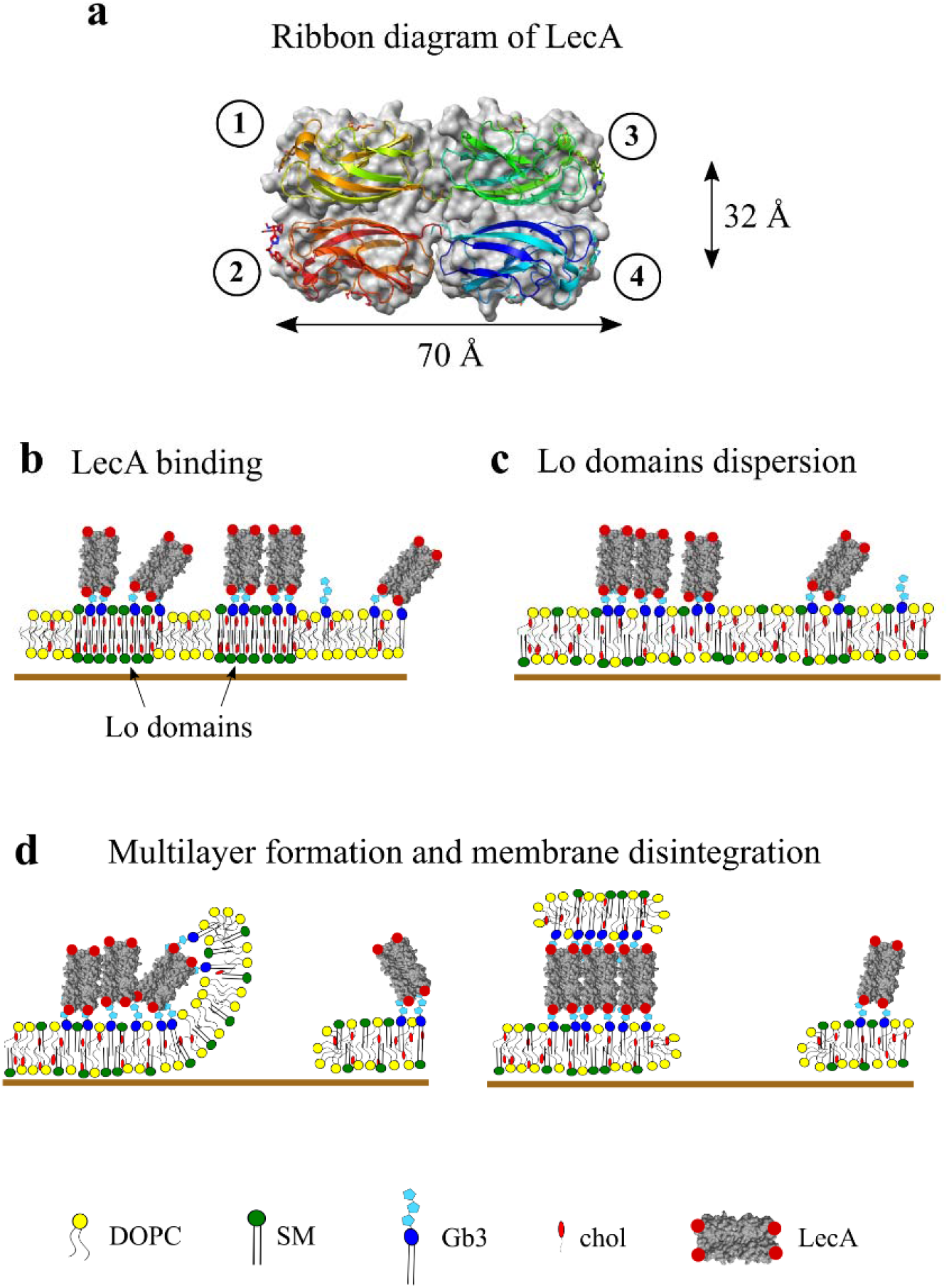
Schematic representation of the membrane reorganization induced by LecA. **a)** Ribbon diagram of LecA with Gb3 binding sites marked by numbers 1-4. **b)** LecA binds Gb3 molecules preferentially in Lo domains. **c)** LecA induces the dispersion of Lo domains. **d)** LecA forms membrane multilayers and induces membrane defects.

### Which factors are important for LecA-induced Lo domains reshaping and dispersion?

LecA and StxB differ in the number and geometry of their Gb3 binding sites. The homo-pentameric StxB contains 15 Gb3 binding sites, which are all oriented in the same direction. LecA has only 4 Gb3 binding sites, with two pairs always pointing in opposing directions (figure 5a). Moreover, the distances between the binding pockets of LecA are larger than in the case of StxB ^22^. This means that LecA cannot be as efficient as StxB in clustering Gb3 molecules, which is crucial for inducing of phase separation (e.g. novel Lo domain formation). However, the configuration of LecA is obviously favorable for inducing the dispersion of Lo domains. Moreover, di-LecA with only two Gb3-binding sites efficiently induces the dispersion of Lo domains.

Next, the binding specificity of LecA to Lo and/or Ld domains was shown to be important for the dispersion of Lo domains. StxB can recognize Gb3 molecules exclusively in Lo domains, whereas LecA is more ambiguous: it binds well to Ld domains as well (figure 5b). Such binding to both Lo and Ld is required for the dispersion of Lo domains, as we demonstrated that both LecA and di-LecA cannot disperse Lo domains when they bind exclusively to Ld domains. Experiments with LecA binding exclusively to Lo domains would provide further insight. Unfortunately, most of Gb3 species yet available demonstrate in the best-case preferential incorporation in Lo domains^41,44-46^. Hereby, we conclude that LecA structure, valence as well as its specificity to membrane domains are decisive factors for the dispersion of Lo domains.

Notably, we previously demonstrated that StxB and LecA trigger different signaling^14^ and intracellular trafficking pathways^24,26^. These differences may originate already at the plasma membrane due to their differences in binding patterns to distinct Gb3 species^41^ and differences in membrane reorganization events they induce. These data in combination with our findings in the current publication indicate the key importance of the geometry and number of their binding sites besides lectin specificity to distinct host cell receptor but also.

### What are membrane multilayers and why do they form?

LecA induces rapidly growing membrane patches on top of the initial SLB. We believe that these membrane multilayers are built from the lipids and membrane fragments of the initial SLB. They consist of two or more stacked lipid bilayers with LecA bound to Gb3 receptors in between. Bright fluorescence signal of LecA associated with such multilayers suggests enrichment and tight clustering of the lectin inside such bilayer “sandwich”. The AFM measurements provided more detailed information on how these membrane multilayers are organized, and moreover, how LecA is oriented when in contact with the membrane. Individual LecA molecules at the initial binding events (height ≈ 2 nm, figure 2) are likely positioned “horizontally” or “tilted” (with its longest axis parallel or at an angle to the lipid bilayer plane). Later, LecA clusters grow (figure 2, height ≈ 9 nm), where LecA is already positioned “vertically” (with its longest axis perpendicular to the lipid bilayer plane). This means that LecA clustering precedes membrane multilayer formation. Furthermore, we suggest that such “vertical” LecA positioning is further preserved in membrane multilayers, with one pair of binding sites facing the initial lipid bilayer at the bottom and with the second pair of binding sites facing the novel lipid bilayer (i.e. top bilayer). Such membrane cross-linking by LecA was observed before in experiments with synthetic GUVs ^39^. Interestingly, membrane multilayer observed by AFM in figure 2 is 24 nm high, which suggests that LecA is also bound on top of the novel lipid bilayer.

The formation of membrane multilayers coincides with the dispersion of Lo domains. Moreover, we found that Lo domains dispersion determines the lipid composition of the membrane multilayers. Membrane multilayers are likely composed out of mixture of lipids from Lo and Ld domains. We concluded this by measuring the membrane order in the multilayers (figure 3).

In homogeneous SLBs as well as in SLBs supplemented with Gb3-FSL (no LecA binding to Lo domains) membrane multilayers formed in absence of the dispersion of Lo domains. Moreover, the formation of membrane multilayers was much faster in such systems.

Dimeric di-LecA did not induce the formation of membrane multilayer, which demonstrates that the geometry of the lectin with two pairs of binding sites facing opposite directions defines the multilayer formation. Similarly, StxB that has as much as 15 binding sites but with all of them facing one direction cannot induce such multilayer structures. Based on our findings, we hypothesize membrane multilayers are formed by cross-linking of the initial SLB “to itself” with subsequent folding of the membrane (figure 5d).

### Can the dispersion of Lo domains occur in absence of multilayer formation?

di-LecA did not induce the formation of multilayers, but it reshaped Lo domains and decreased their total areas. Moreover, it induced an apparent asymmetric mixing of lipids, by fully dispersing Lo domains only in the top leaflet of the SLB, while the Lo domains in the bottom leaflet remained intact. Hence, the dimeric geometry of di-LecA is sufficient for the dispersion of the Lo domains, whereas the native tetrameric geometry of LecA with two pairs of binding sites facing opposite directions is absolutely necessary for the formation of membrane multilayers.

### What might be the physiological role of LecA-induced Lo dispersion and multilayer formation?

In nature, LecA is involved in the invasion by *P. aeruginosa* of the host cells via the lipid zipper mechanism^11^. Moreover, it is crucial for bacterial biofilm formation^47^. The lipid zipper is formed by attachment of *P. aeruginosa* to the host cell by LecA. Two LecA binding sites are probably attached to the bacterial surface, whereas the two other binding sites probably bind to Gb3 molecules in host cell membrane. Similar bacterium-host cell surface as well as bacterium-bacterium association mediates the formation of biofilms. Here, we demonstrated that LecA can not only cross-link already existing membranes but also cross-link membrane “to itself” and form micrometer-sized membrane multilayers which shows a very peculiar mechanism of membrane deformation that can be explained as “folding” or “wrapping”. This previously unknown feature can be utilized by bacteria during initial steps of invasion for membrane deformation.

Furthermore, LecA disperses Lo domains in the membrane lateral plane. Tetrameric as well as dimeric LecA can do it, which means that when LecA is attached to the bacterial surface and exposed to the host cell membrane only with two binding sites, it still can disperse ordered domains. Several publications demonstrated that host cell infection of *P. aeruginosa* is mediated by interaction with ordered nanodomains (also mentioned before as lipid rafts) in the plasma membrane^48–51^. The dispersion of Lo domains showed here provides more mechanistic insight into the impact of lectins and *P. aeruginosa* on ordered domains. Such membrane reorganization dynamics can be interpreted as a prerequisite for induction of signaling cascades as well as membrane mechanical modification (e.g. bending rigidity decrease) to facilitate plasma membrane deformation.

## Conclusion

In this work, we found that the lectin LecA from *P. aeruginosa* induces the dispersion of ordered domains in model membranes. These findings provide new insights into the mechanism of *P. aeruginosa* invasion in host cells as the dispersion of membrane ordered domains may be beneficial for lowering membrane bending rigidity and favoring the lipid zipper and bacterial internalization. Furthermore, we compared the impact of two Gb3-binding lectins, LecA and StcB on model membrane organization. We found that not only specificity but also lectin geometry determines the lectin behaviour on the membrane surface.

## Acknowledgments

This study was supported by the German Research Foundation (DFG) under Germany’s Excellence Strategy (CIBSS – EXC-2189 – Project ID 390939984, BIOSS – EXC 294), by DFG grants RO 4341/3-1, Major Research Instrumentation (project numbers: 438033605 and 290424854), RTG 2202 (278002225) and IRTG 1642, by the Ministry of Science, Research and the Arts of Baden-Württemberg (Az: 33-7532.20), by the German Federal Ministry of Education and Research (BMBF) in the framework of the EU ERASynBio project SynGlycTis, by the Freiburg Institute for Advanced Studies (FRIAS), and by a starting grant from the European Research Council (Programme “Ideas,” ERC-2011-StG 282105). T.Sy. acknowledges support by the Franco-German University (programs ‘Polymer Sciences’ and ‘Cotutelle de thèse’) and the Collège Doctoral Européen (PDI). Y.M. is grateful to the Institut Universitaire de France (IUF) for support and providing additional time to be dedicated to research. We are grateful to Andrey Klymchenko for providing the lipophilic membrane probes.

## Notes

### Competing Interest Statement

The authors have declared no competing interest.

