## Supplementary figures for "The bacterial lectin LecA from *P. aeruginosa* alters membrane organization by dispersing ordered domains"

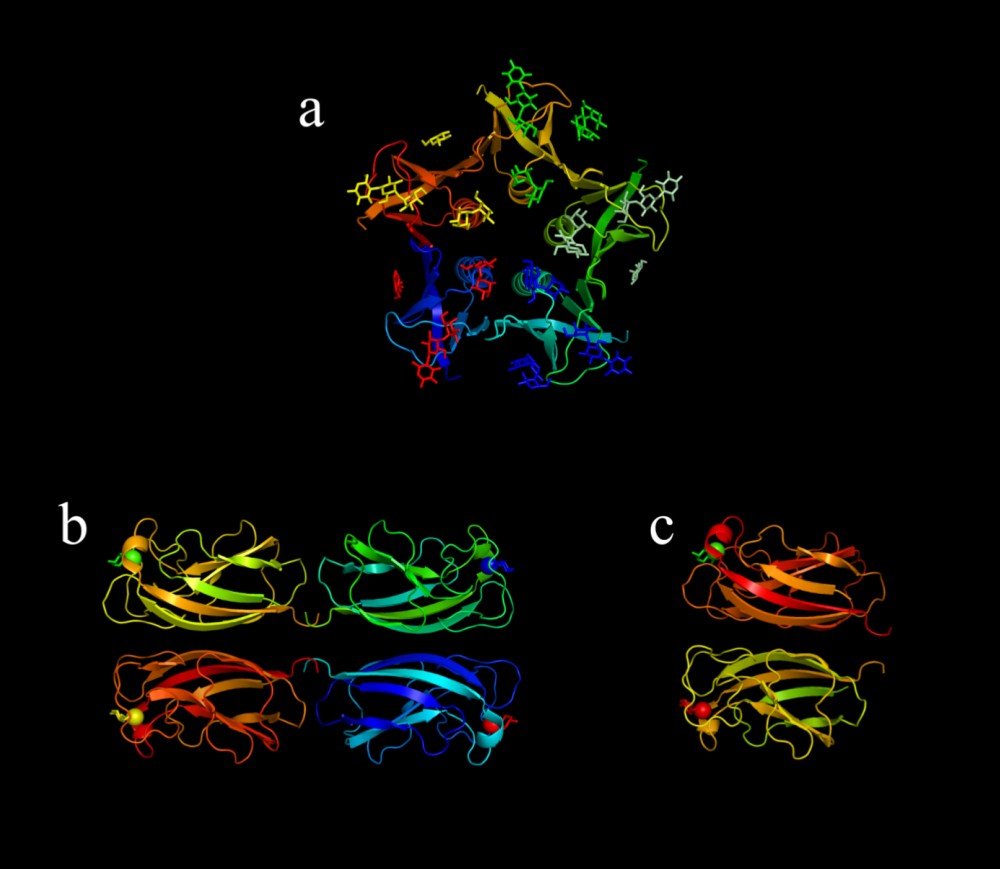

Figure S1|**Ribbon diagrams of lectins.** The Gb3 binding sites are marked by the carbohydrates and Ca^2+^ ions. **a)** Shiga toxin B subunit (StxB) from *S. dysenteriae*; Adapted from protein data bank (PDB, 2xsc) **b)** LecA from *P. aeruginosa*; Adapted from protein data bank (PDB, 4cp9) **c)** Recombinant prokaryotic lectin $\alpha$ Gal (RPL, further referred to as di-LecA). Adapted from protein data bank (PDB, 4cp9). The structures in a and b were provided by RCSB Protein Data Bank and visualized by Pymol software. The ribbon diagram in c is presumptive.

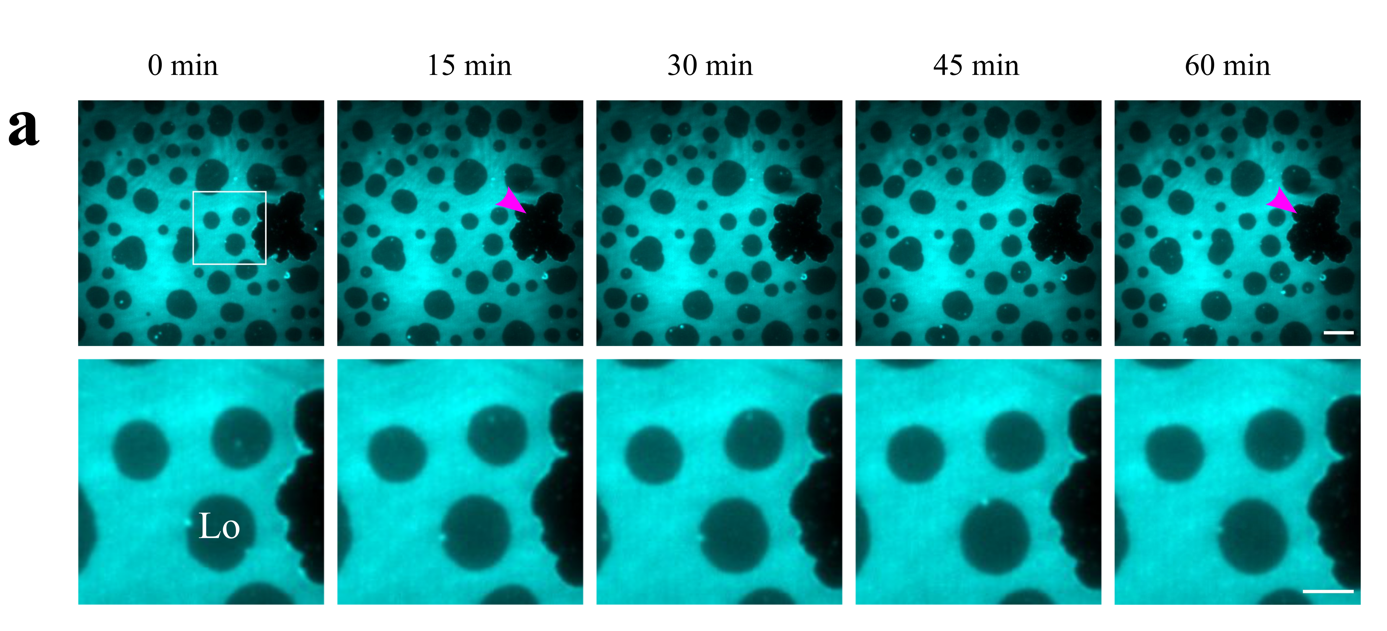

Figure S2| **Negative control.** Phase separated SLB labeled by HPC-Bodipy was imaged during one hour using widefield microscopy (HILO mode). The Lo domains and their lateral distribution are stable during one hour of acquisition. A membrane defect (magenta arrowhead) that formed during SLB preparation is stable and neither expands nor shrinks during the acquisition. Scale bars – 10 and 5 µm respectively.

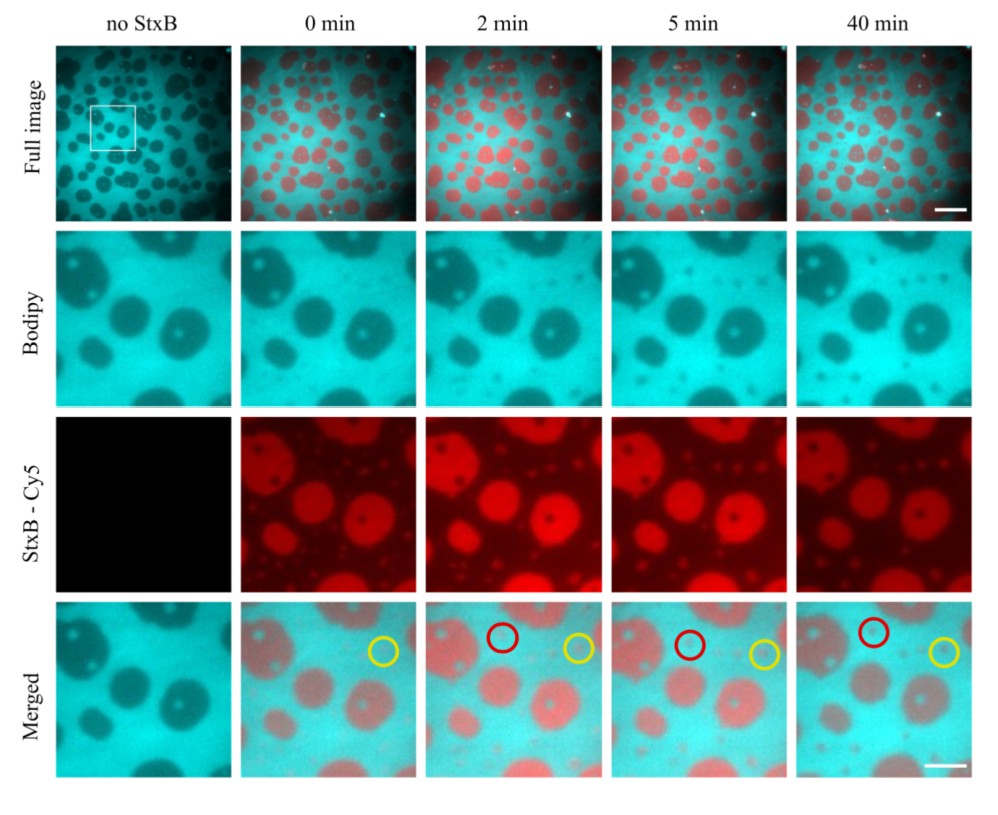

Figure S3| **StxB-induced SLB reorganization.** The SLB was composed of DOPC/chol/SM/Gb3 (37.5/20/37.5/5). The SLB was labeled by Bodipy-HPC that preferentially incorporates in Ld domains. StxB-Cy5 (200 nM) was applied. StxB binds almost exclusively to Lo domains. Moreover, it induces a formation of new Lo domains (red and yellow circles - Merged) by efficient clustering of the Gb3 molecules incorporated in the Ld phase. Scale bars – 10 and 5 µm respectively.

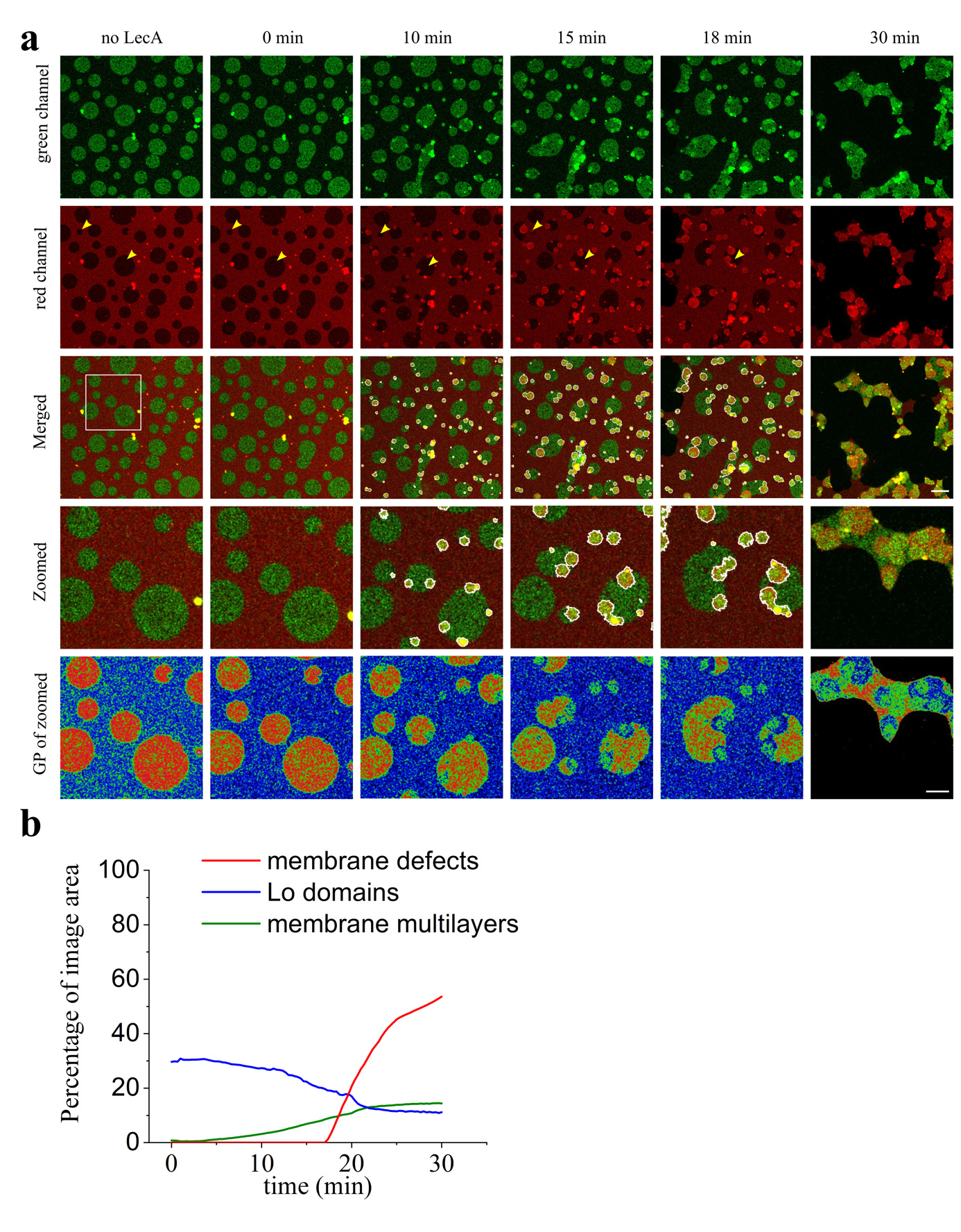

Figure S4|**Time series of the two-color imaging for membrane order mapping.** SLB is labeled by NR12S. LecA is unlabeled. Green channel is acquired using the 525/50 BP emission filter, red channel is acquired using 700/75 BP emission filter. **a)** The dynamics of the membrane reorganization are in line with the observations presented in main figure 2. Lo domains decrease in size and finally disappear (yellow arrowheads). Membrane multilayers form, they are depicted as ROIs with white borders. **b)** Total areas of membrane defects, Lo domains, Ld domains and membrane multilayers over time. Scale bars – 10 µm and 5 µm respectively. The complete time sequence is available as the supplementary movie 2.

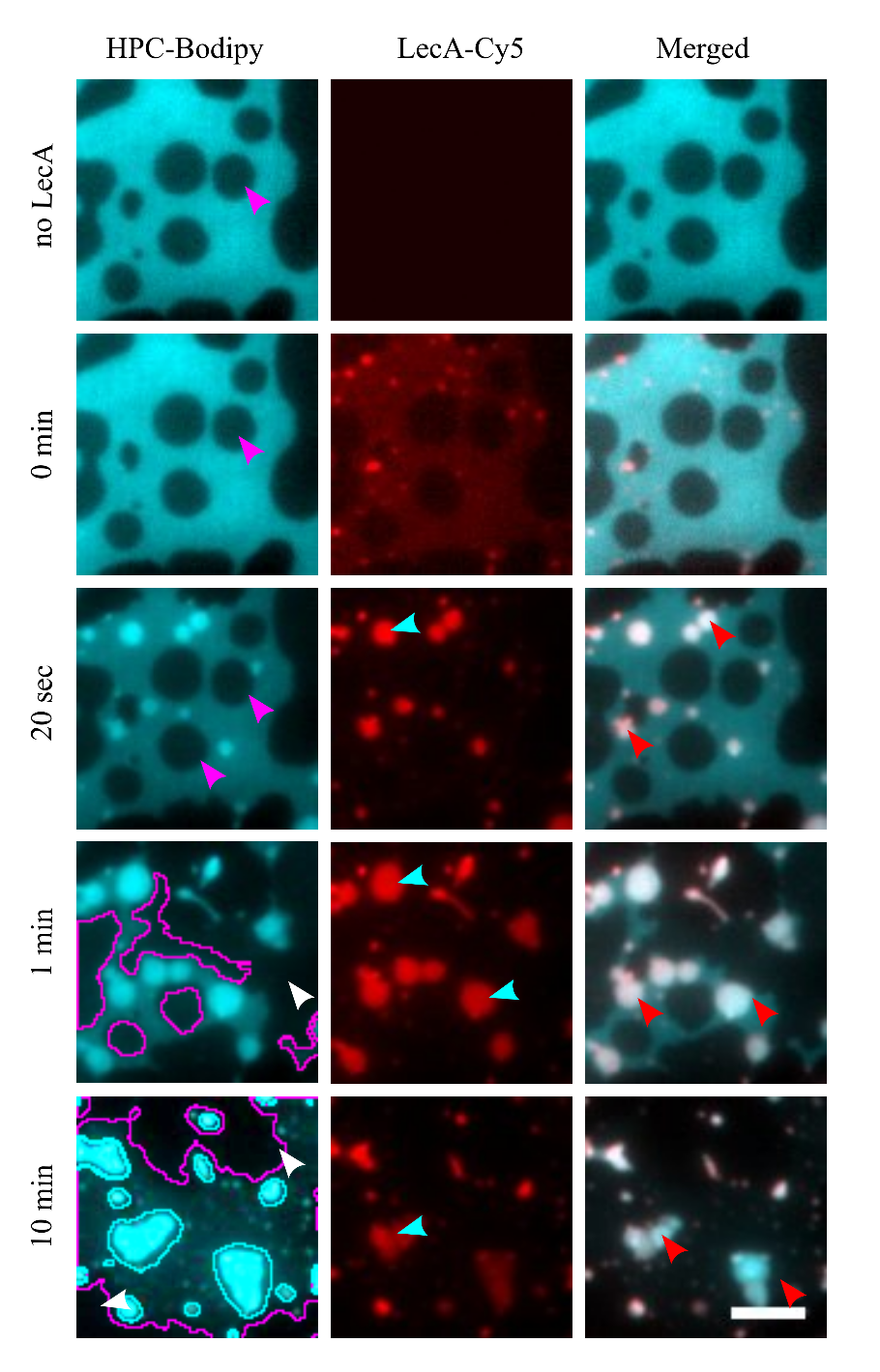

Figure S5| **LecA-induced reorganization of SLBs supplemented with Gb3 FSL.** Time series of LecA interaction with phase-separated SLB (DOPC/chol/SM (37.5/20/37.5)) supplemented with 5 mol % of Gb3-FSL. SLBs are labeled by HPC-Bodipy that localizes to Ld domains (cyan). LecA is labeled with Cy5 (red). Images are acquired with wide field microscopy (HILO). In SLB with Gb3 FSL, LecA binds preferentially to Ld. LecA induces Lo domains reshaping, but no dispersion (white arrowheads). LecA clusters at the SLB (cyan arrowhead) and induces the formation of membrane multilayers (red arrowhead). Already after 10 min membrane defects occur (purple arrowhead). At 1 min and 10 min time points, Lo domains are highlighted with magenta ROI and membrane defects are indicated with white arrowheads. Membrane multilayers are encircled with cyan ROIs. Scale bar – 5 µm. The complete time sequence is available as the supplementary movie 4.

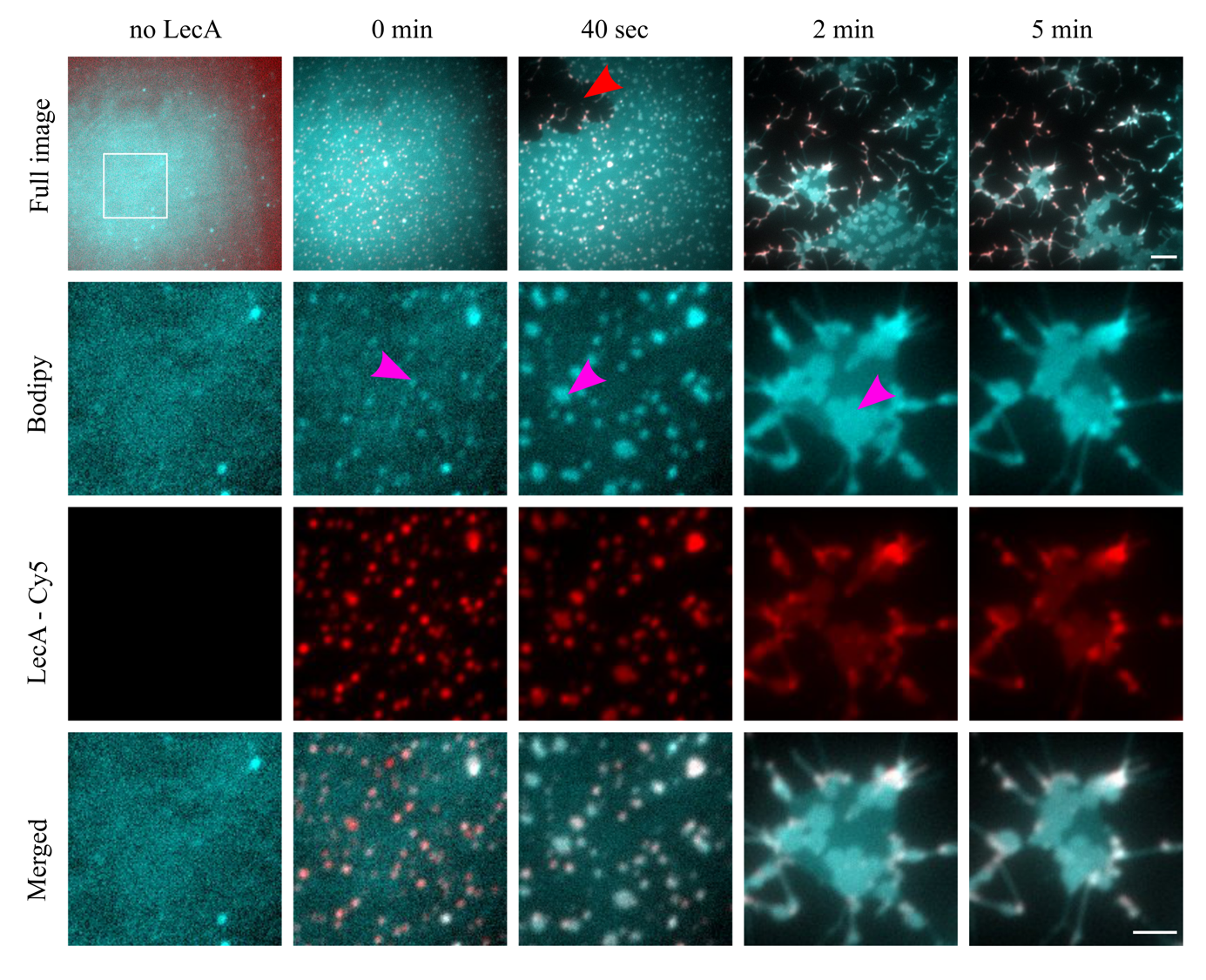

Figure S6 | **LecA-induced membrane reorganization of homogeneous SLB.**  The SLB was prepared from a non-phase separating lipid mixture (DOPC/chol/Gb3 (65/30/5). Membrane marker – HPC-Bodipy). Since such lipid bilayers do not exhibit phase separation, the membrane fluorescence signal (cyan) appears homogeneous. Application of LecA labeled with Cy5 (red) induces membrane multilayers formation (magenta arrowheads) and fast membrane disintegration (membrane defect is highlighted with red arrowhead). Scale bars – 10 and 5 µm respectively. The complete time sequence is available as the supplementary movie 3.

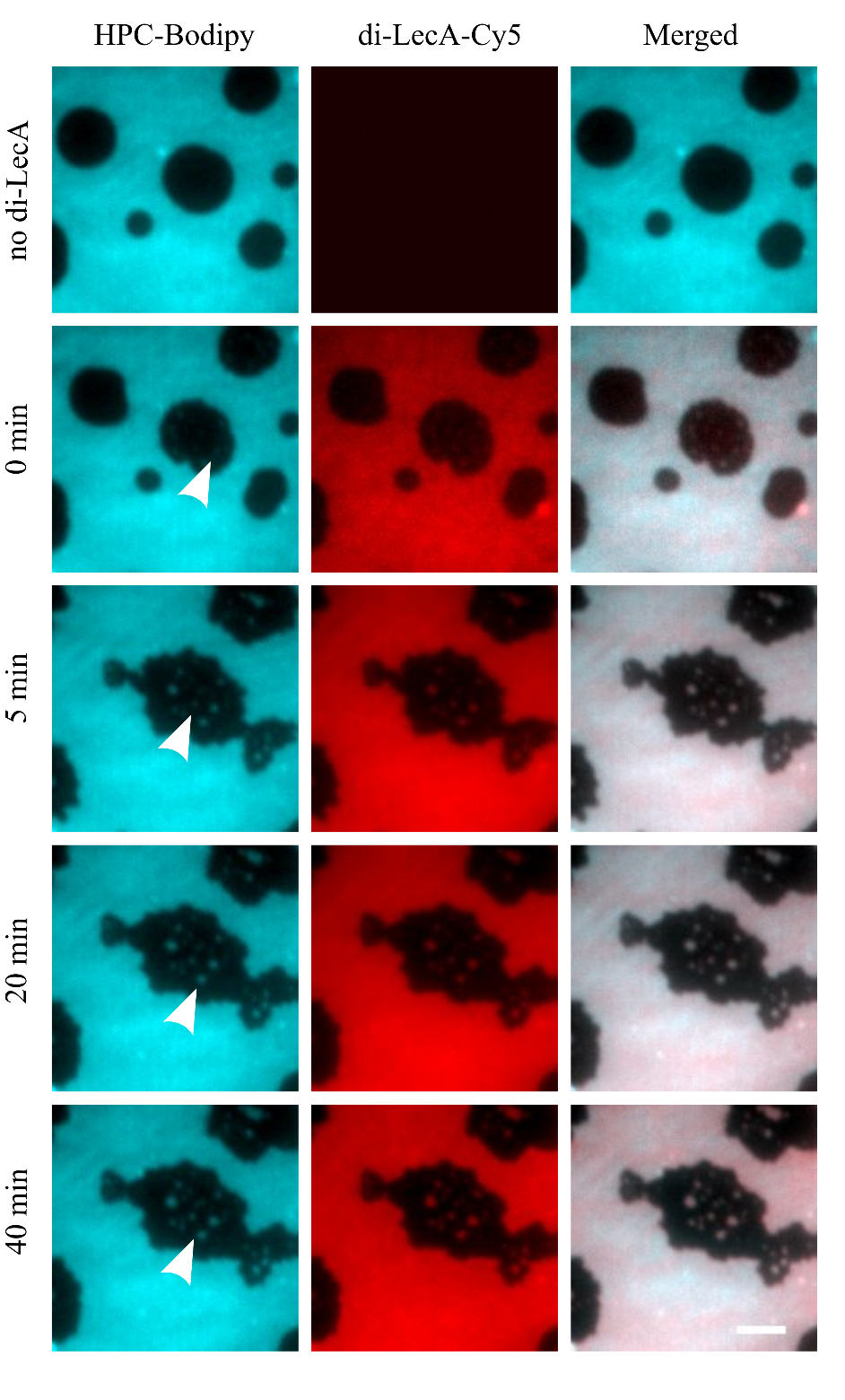

Figure S7| **Di-LecA-induced reorganization of SLBs supplemented with Gb3 FSL.** Time series of di-LecA interaction with phase-separated SLB (DOPC/chol/SM (37.5/20/37.5)) supplemented with 5 mol % of Gb3-FSL. SLBs are labeled by HPC-Bodipy that localizes to Ld domains (cyan). RPL is labeled with Cy5 (red). Images are acquired with wide field microscopy (HILO). In SLB with Gb3 FSL, di-LecA binds preferentially to Ld. di-LecA does not induce Lo domains dispersion. However, Lo domains are re-shaping and fusing (white arrowhead); Scale bar – 5 µm. The complete time sequence is available as the supplementary movie 6.

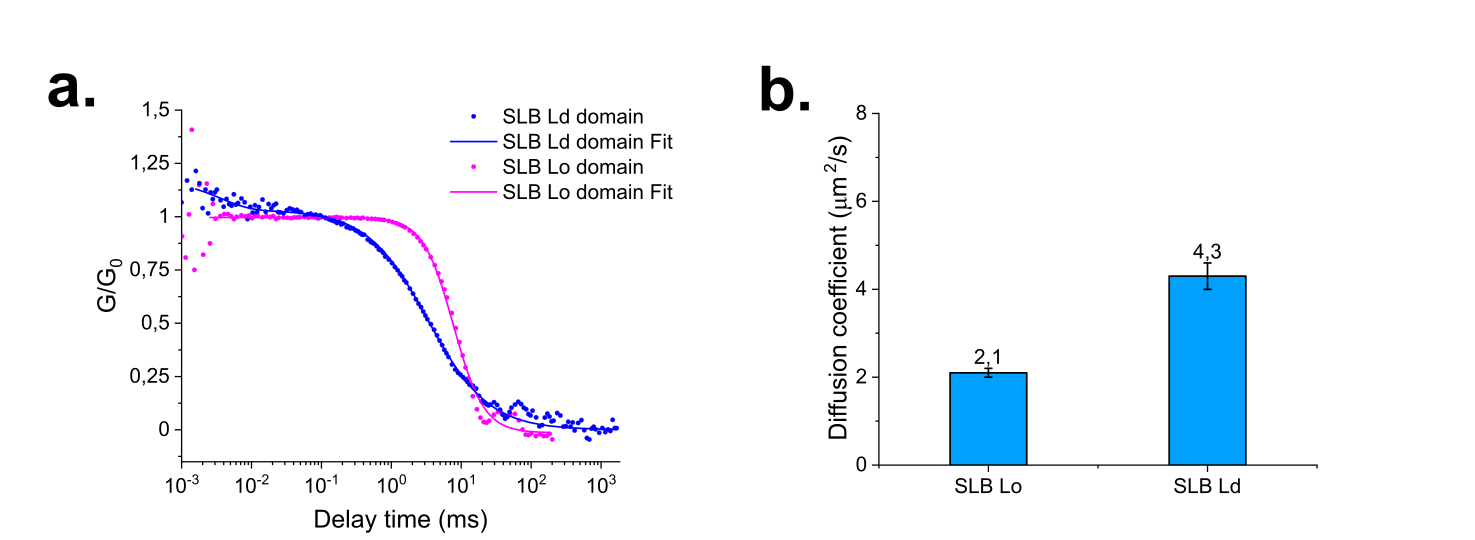

Figure S8| **Mobility control of SLBs using Fluorescence Correlation Spectroscopy.** a) Autocorrelation curves of HPC-Bodipy in Ld and Lo of SLBs on mica (DOPC/SM/Chol/Gb3, 40/40/15/5); b) Diffusion coefficients of HPC-Bodipy in Ld an Lo.

**Supplementary movie 1| LecA induces reorganization of the SLB.** The complete time sequence of the experiment presented in figure 2a and figure S2. LecA binds to the Lo domains and dissolves them (purple arrowheads). The membrane multilayers, induced by LecA are depicted with the white arrowheads. The subsequent formation of membrane defects is depicted by orange arrowhead. Several events indicate possible asymmetric Lo domains dissolution (red arrowheads), similar to the one presented in supplementary movie 5.

**Supplementary movie 2| Ratiometric imaging of the LecA-induced SLB reorganization.** The complete time sequence of the experiment presented in figure 2 and figure S3. The dissolving of the Lo domains (one of many events is pointed by white arrowhead) and multilayers formation (two of many events are pointed with cyan arrowheads) are depicted here.

**Supplementary movie 3| The reorganization of the homogeneous SLB induced by LecA.** The complete time sequence of the experiment presented in figure S4. The phase separation is absent in this SLB. LecA binds to the SLB and induces the multilayers formation (white arrowheads). Subsequently, the membrane defects are induced (orange arrowhead).

**Supplementary movie 4 | The LecA-induce reorganization of the SLB supplemented with Gb3 FSL.** The movie depicts the complete time sequence of the experiment presented in figure 3. The reorganization of the Lo domains is depicted by white arrowheads and the multilayer is depicted with blue arrowheads.

**Supplementary movie 5| Di-LecA induces reorganization of the SLB.** The complete time sequence of the experiment presented in figure 4a. Di-LecA binds to the Lo domains and induces their partial dissolution (white arrowheads). Thereafter, the re-shaping of the Lo domains stops, but they are dissolved asymmetrically.

**Supplementary movie 6 |The di-LecA-induced reorganization of the SLB supplemented with Gb3 FSL.** The complete time sequence of the experiment presented in figure 5. The reorganization of the Lo domains is depicted by white arrowheads.
